# Visual Cortical Area MT is Required for Development of the Dorsal Stream and Associated Visuomotor Behaviours

**DOI:** 10.1101/2021.02.28.433301

**Authors:** William C. Kwan, Chia-Kang Chang, Hsin-Hao Yu, Inaki C. Mundinano, Dylan M. Fox, Jihane Homman-Ludiye, James A. Bourne

**Author notes:** Authors contributed equally (William C. Kwan, Chia-Kang Chang).

## Abstract

The middle temporal (MT) area of the extrastriate visual cortex has long been studied in adulthood for its distinctive physiological properties and function as a part of the dorsal stream, yet interestingly possesses a similar maturation profile as the primary visual cortex (V1). Here we examined whether an early-life lesion of MT altered the dorsal stream development and the behavioural precision of reaching to grasp sequences. We observed permanent changes in the anatomy of cortices associated with both reaching (PE and MIP) and grasping (AIP), as well as in reaching and grasping behaviours. In addition, we observed a significant impact on the anatomy of V1 and the direction sensitivity of V1 neurons in the lesion projection zone. These findings indicate that area MT is a crucial node for the development of the primate vision, impacting both V1 and areas in the dorsal visual pathway known to mediate visually guided manual behaviours.

**Teaser:** The early life loss of visual area MT leads to significant anatomical, physiological and behavioural changes.

## Introduction

Vision relies on a multiplicity of areas within the brain to communicate synergistically in order to produce an accurate percept. The seminal works of Goodale & Milner(*1*) and Ungerleider & Mishkin(*2*) proposed the existence of two neocortical pathways which have a dichotomous visual functional specialisation. More specifically, the dorsal stream pathway is involved in perceiving motion and “vision-for-action”, while the ventral stream pathway is associated with shape/ object perception and “vision-for-perception”. Central to the dorsal stream is the middle temporal (MT) area **(Fig. 1A, B)**. Based on established anatomical markers of neural circuit maturation, area MT matures early and in parallel with primary sensory areas, including the primary visual cortex (V1)(*3*). Further, since the monosynaptic V1-MT connection is yet to be fully established during this process, the early maturation of MT is thought to be independent of V1 input(*4*). This knowledge, and the fact that MT has a first-order topographical map(*5*) and is directly recipient of thalamic connections(*6*, *7*), supports the notion that it is a ‘primary-like’ area and serves as an anchor early in life to support establishment of the dorsal stream(*8*).

**Figure 1.**
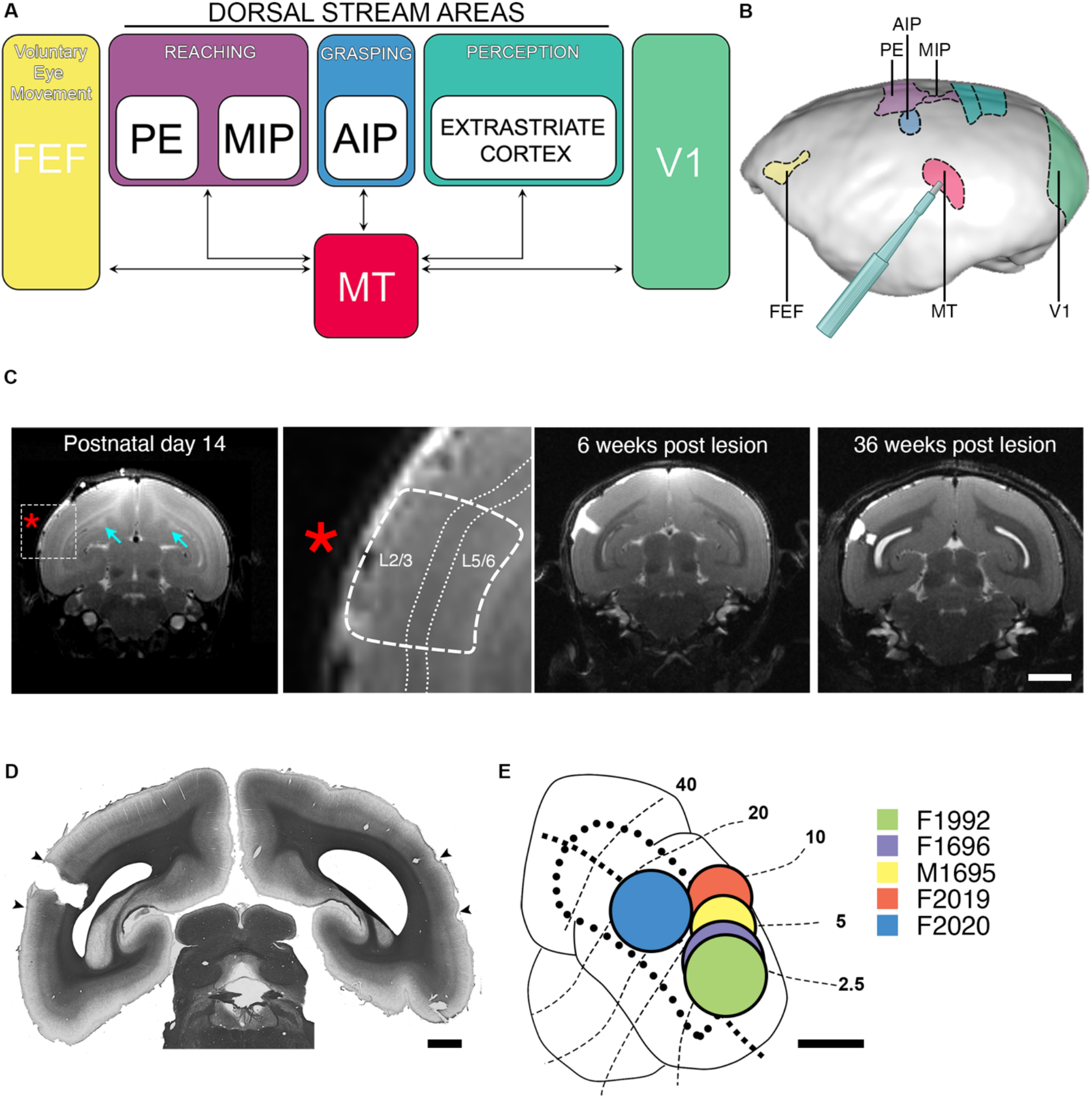
Direct connectivity of middle temporal (MT) area of the visual cortex and unilateral early life lesion approach. **(A)** Wiring diagram detailing monosynaptic reciprocal connectivity of area MT and proposed behavioural function. (**B)** Lateral view of left hemisphere of marmoset cortical surface highlighting anatomical positioning of areas in (**A)**. (**C)** T2-weighted MRI of a marmoset brain at the level of parafoveal MT, which was targeted for ablation at PD14. MT was demarcated based on dorsal shift in the layer IV (left image, red asterisk and inset). In the anterior/posterior plane, the core of MT was denoted as when the most caudal aspect of the corpus callosum becomes visible (arrow). Further T2-weighted scans were conducted at 6 weeks and 1 year post-injury to confirm lesion location and also diffusion tractography imaging (DTI) for fractional anisoptropy analysis (DTI results in **Fig. 4**, scale bar = 5mm). **(D)** Histological analysis with myelin silver staining for *ex vivo* demarcation of lesion (scale bar = 2mm) **(E)** Summary of the extent, and retinotopic relationship, of MT lesion in each lesioned animal (scale bar = 2mm). Details of the experimentation each animal underwent is further detailed in **Table 1**.

In adulthood, MT receives significant input directly and indirectly from V1(*5*, *9*), and is interconnected with multiple dorsal stream extrastriate areas, including V3 and the dorsomedial (DM) area, anterior and lateral intraparietal areas (AIP and LIP) (*10*–*13*) and the frontal eye fields (FEF)(*14*, *15*) **(Fig. 1A, B)**. Area MT also has abundant connections with subcortical structures, including reciprocal connections with the medial portion of the inferior pulvinar(*4*, *7*, *16*), and afferent projections from the koniocellular layers of the lateral geniculate nucleus (LGN) (*6*) and superior colliculus (*11*, *17*, *18*).

A substantial proportion of neurons within MT are highly tuned for direction(*19*, *20*), a clear distinction from V1, suggesting that MT plays a central role in motion perception and integration of visual information. However, there are few studies which have interrogated how injury to MT impairs visual perception and global visual behaviour. Newsome and colleagues(*21*, *22*) provided behavioural evidence of how MT contributes to motion perception following mechanical ablation of MT, while motion blindness/ akinetopsia has been observed clinically in a patient with bilateral MT injury(*23*, *24*). Further, others have demonstrated that V2 direction tuning properties are dependent on MT(*25*).

In the phenomenon of blindsight, the extensive multiplexed circuitry of the dorsal stream, with a central component being area MT, has been long implicated as the neural substrate that affords the ability to perform specific visually-driven tasks in the absence of V1(*26*, *27*). More specific investigation revealed that MT remains active when visual stimuli are presented within the scotoma(*28*, *29*).

Few studies have considered the role of MT in the development of the dorsal stream and to the establishment of specific visuomotor behaviour but developmental studies have inferred that MT acts as the primary node the establishment of the dorsal stream(*3*, *30*). Further, early-life lesions of the geniculostriate pathway have highlighted the conserved integrity of MT in both humans and monkeys (*27*, *31*). Therefore, we hypothesised that an early-life lesion of MT will permanently perturb reaching to grasp behaviours and result in dysfunction of areas directly connected with MT.

## Results

We used early postnatal ablations as a method to study the role MT would serve in the development of the visual cortex and visually-guided behaviours. At 6 weeks post-surgery, MRI T2-weighted images were acquired to validate unilateral biopsy punch ablation of MT in the 5 neonatal (PD14) marmosets **(Fig. 1C)**. Prior to lesion, the location of area MT was visualised in MRI images (T2-weighted) by a dorsal shift in layer 4 from adjacent areas **(Fig. 1C inset)**. During the excision of MT tissue, particular care was taken to remove all 6 cortical layers while leaving the underlying white matter tract intact. When the animals subsequently reached young adulthood (>36 weeks) and underwent DTI scans for evaluation of effected cortices, another T2 sequence was obtained to map the extent of the lesion **(Fig. 1B)**. In these scans, we often observed that the lesion core had scar tissue and subtle degeneration of the underlying white matter had occurred. Following perfusion of the cerebral tissues, histological confirmation of the lesion was achieved with a Gallyas (myelin) silver stain **(Fig. 1D)**, and the lesion was fully reconstructed to establish its topographical extent for each animal **(Fig. 1E)**.

### The absence of MT during development perturbs reach and grasp kinematics as well as interaction with static and moving objects

As MT neurons are highly tuned for direction and the perception of motion, we wanted to determine if accurate reward-driven visuomotor behaviour was still achievable following early-life loss of MT. To ensure adequate compliance with the task, we first trained the animals to enter a removable transport box on the home cage, which was then taken into the behaviour laboratory **(Fig. 2A)**(*32*). We employed naturalistic, goal orientated, reach and grasp tasks once neonatal MT lesion animals (PD14) had reached adulthood (>18 months). These were undertaken both statically **(Fig. 2B)** or upon a variable speed/ direction turntable **(Fig. 2b)**. Each food retrieval trial was 2D video-captured (see camera; **Fig. 2A**), which allowed us to study the animal’s proficiency in executing the task and the kinematics of the reach and grasp phases.

**Figure 2.**
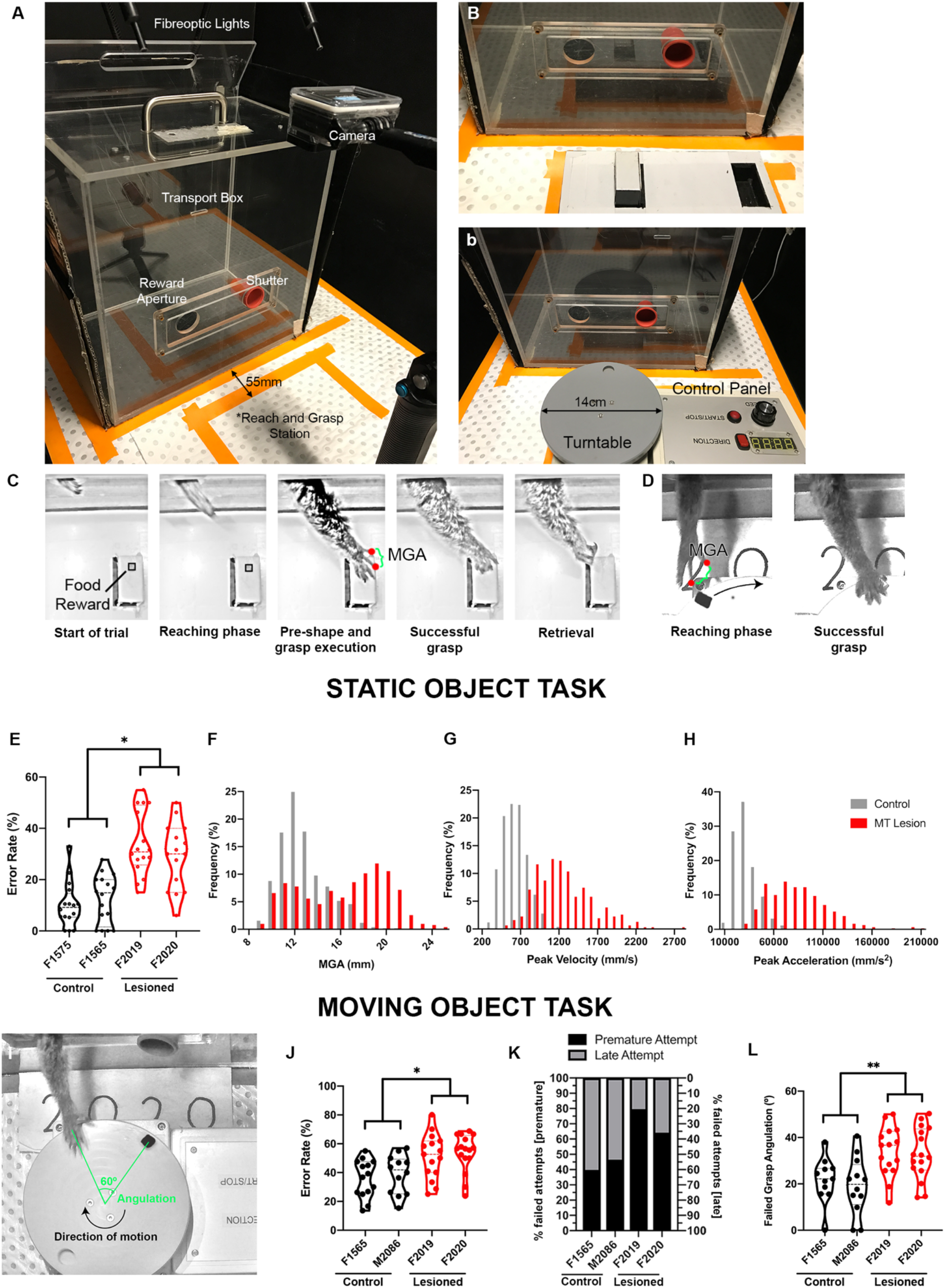
Early-life MT lesions perturb goal orientated reach and grasp performance in static and moving objects. **(A)** Experimental setup of reach and grasp tasks. The animal is presented with either a static task **(B)** or moving task **(b)** (see methods for full details). **(C)** Example of 720p footage of an animal successfully grasping a static reward. **(D)** Example of 720p footage of an animal successfully grasping a moving reward. **(E)** Performance (error rate) of control vs lesioned animals in reaching and grasping static objects. Lesioned animals performed significantly worse (permutation test, *p*=0.04*). **(F)** Frequency distribution across trials of maximum grip aperture (MGA) for control and lesioned animals. Lesioned animals exhibit a larger MGA in their grasp and reach with greater velocity **(G)** and acceleration **(H)**. **(I)** An example trial where the animal was delayed in reaching for the food reward and failed in the moving object reach and grasp trial, the magnitude of which was calculated. **(J)** Performance of control vs lesioned animals in reaching and grasping moving objects. Lesioned animals performed significantly worse (permutation test, *p*=0.042). **(K)** Failed trials within the lesioned cohort were largely due to the animal reaching prematurely. Control animals showed close to a 1:1 proportion of premature and late reaching actions in failed trials whilst MT lesioned animals exhibited a tendency to reach prematurely. **(L)** The average magnitude of error as denoted by angulation in the failed trials. When lesioned animals failed in reaching and grasping moving objects, the magnitude of the miss was greater than controls (permutation test, p=0.003).

Animals were first presented with a static task. The task involved presenting a food reward which was placed on a pedestal in front of the animal **(Fig 2B - D)**. Food rewards were cut into two sizes of cuboid (*small*: ~5×5 mm face; and, *large*: ~10×10 mm face) and were first employed to determine if the object size would affect overall performance. No significant difference in retrieval performance was observed between the two sizes of fruit reward in both control and lesioned cohorts (ANCOVA F(1,52)=0.68, *p*=0.41)(Fig S1A, B). When examining the collective retrieval attempts of both sizes of the statically presented food reward, we found overall performance within the MT-lesioned group was significantly impaired when compared with controls (permutation test, *p*=0.04*)**(Fig. 2E)**. Further analysis of the reach and grasp kinematics revealed significant differences in hand preshaping and reaching behaviour. Specifically, during the preshaping phase, neonatal MT-lesioned animals exhibited a significantly larger maximum grip aperture than controls (permutation test, *p*=0.0037) **(Fig. 2F)**. Further characterisation of the behaviour revealed neonatal MT-lesioned animals exhibit a feedforward movement with greater velocity (permutation test, *p*=0.04) **(Fig. 2G)** and acceleration (permutation test, *p*=0.02)**(Fig. 2H)** when approaching the target object.

We further assessed whether a target object of different sizes may influence the animal’s ability to pre-shape their hand. By plotting the object size against the MGA, a significant correlation was observed between the two cuboid food pieces in the control group (Pearson’s correlation, r(381)=0.16, *p<*0.01**)(Fig S1C). However, hand preshaping is not seen within the neonatal MT-lesioned group (Pearson’s correlation, r(338)=0.05, *p*=0.33)(Fig S1D).

To interrogate the animal’s ability to accurately engage a moving object in a naturalistic setting, animals were again required to reach and grasp a food reward, but in this instance the reward was placed on a variable speed/ direction turntable in front of the transport box **(Fig. 2b, I)**. A difference in object retrieval error rate was observed between the two cohorts, with the neonatal MT-lesioned group having an increased error rate (permutation test, *p*=0.042) **(Fig. 2J).** Further analysis of reach and grasp behaviour within the lesion cohort revealed that when retrieval of the food reward was unsuccessful, this was largely due to executing their reaching action prematurely **(Fig. 2K)**. When lesioned animals failed in retrieval of moving objects, the magnitude of their failure was greater compared to controls (permutation test, p= 0.003)**(Fig. 2L)**. Further, they were less co-ordinated and consequently had less success in retrieval of the reward on their first attempt (permutation test, *p*=0.023)(Fig. S1E). The MGA, peak velocity and acceleration observed between both cohorts for the moving task were similar (Fig. S1F, G, H).

### V1 responses are altered following an early-life lesion of MT

As there is a significant monosynaptic projection between V1 and MT, we wanted to observe if an early-life lesion of MT impacted upon the response properties of neurons in the lesion projection zone (LPZ) of ipsilesional V1 in adulthood. Visually evoked single-unit activity was recorded in the LPZ of V1 from two animals (M1695 & F1696, **Fig. 3A**), with a total of 122 units recorded from. Visual stimuli were presented to determine the orientation and direction selectivity as well as optimal spatial and temporal frequencies for each of the units.

**Figure 3.**
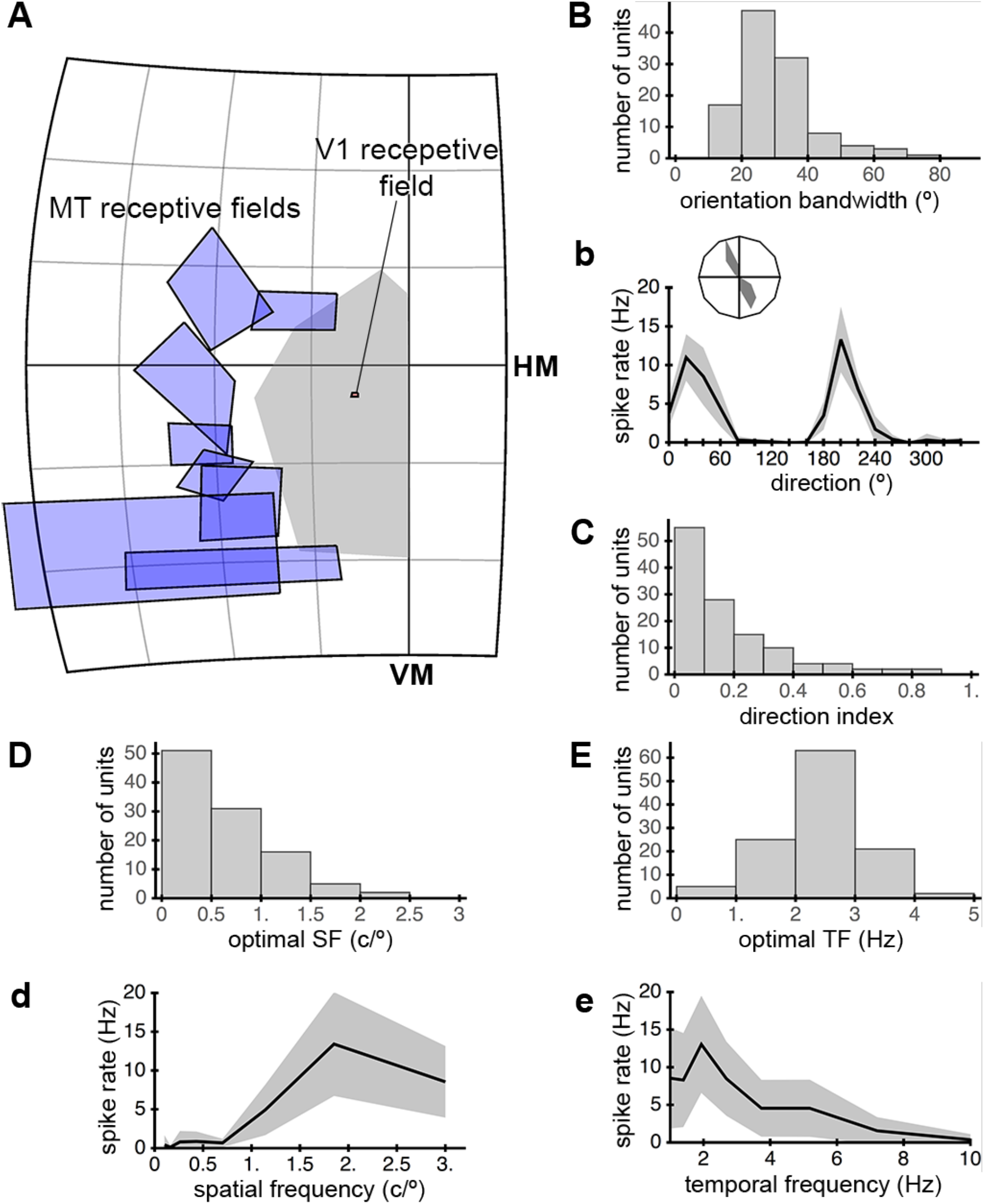
Tuning properties of V1 neurons following unilateral MT lesion. **(A)** MT receptive fields were mapped by recording on the periphery of the lesion core. Following receptive field mapping, the scotoma is delineated and single unit recordings of V1 neurons were performed within the MT-V1 lesion projection zone (LPZ) (see methods for full details). **(B)** The orientation selectivity of all recorded V1 neurons as determined by the half-width-half-height bandwidth (Median HWHH = 29.5° from 122 units). **(b)** An example of the orientation tuning curve of a V1 neuron within the LPZ. **(C)** Direction index of the 122 V1 units recorded within the LPZ. A direction index of >0.5 suggests the neuron is directionally selectivity. Only 10 out of 122 (~8%) of neurons were tuned for direction. V1 neurons within the LPZ responded optimally to a spatial frequency of 0.7 cycles/° **(D)** and temporal frequency of 2.3 Hz **(E)**. Examples of the spatial **(d)** and temporal frequency **(e)** tuning curves from V1 neuron within the LPZ are shown.

Our first observation was that the basic retinotopic organisation of V1 was unaffected by the neonatal V1 lesion **(Fig. 3A)**, which was expected. Our single-unit recordings revealed that 86.1% (105 out of 122) of V1 neurons within the LPZ were found to be orientation selective. The half-width-half-height (HWHH) bandwidth which is a traditional measure for orientation selectivity(*33*, *34*) within the MT lesion affected V1 was observed to be 29.5° **(Fig. 3B)**. This is comparable to the median HWHH bandwidth of 29.0° which has been reported previously for normal V1 neurons throughout the entire visual field(*35*). Direction selectivity (DS) of a cell was determined by calculating a direction selectivity index(*35*, *36*), whereby an index of >0.5 suggests the cell is tuned for direction. Previous reports within marmoset V1 demonstrate approximately 20% of V1 neurons are DS. Our findings were lower than this, with less than 10% (10 out of 122) classified to be direction selective (DS) **(Fig. 3C)**. Although the proportion of DS can vary with eccentricity: central vision, 18% DS; near periphery, 26% DS; and, far periphery, 20% DS(*35*), our observations of ~10% fell below this previously reported proportion.

Neurons in V1 within the two subjects responded optimally to a spatial frequency of 0.7 cycles/° **(Fig. 3D)** and a temporal frequency 2.3 Hz **(Fig. 3E)**. Previously determined values for optimal spatial frequency was 1.08 cycles/° centrally and 0.45 cycles/° in the near periphery, with optimal temporal frequency between 3.0-4.0 Hz across the entire visual field. Therefore, the spatiotemporal sensitivities appeared unaffected by an early-life MT ablation.

Together, the basic retinotopic layout and physiological properties of V1 neurons appeared unaffected by the early life MT lesion, with one exception. Namely, across the recorded areas, the proportion of directionally selective neurons was lower than previously reported, suggesting that MT may play a role in shaping these responses.

### Early-life lesions of MT lead to widespread changes in cortical architecture

An advantage of our early-life lesion model is that it allows us to perform longitudinal MR imaging following the lesion. We performed whole brain DTI scans on our lesioned animals at 36 weeks to determine the extent of chronic disruption to the cortical architecture. Voxel based morphometric (VBM) analysis of the DTI maps between the lesioned and non-lesioned hemisphere revealed, a reduction in fractional anisotropy (FA) in the intraparietal cortical areas AIP, MIP, posterior parietal area PE (dorsal stream areas), the frontal eye fields (FEFs) and V1 (p value <0.05)**(Fig. 1A; Fig. 4A, B)**, areas MT has strong monosynaptic connections with. This result is suggestive of an altered state in the neuroanatomy of these areas, where a reduction in FA, has been correlated with both axonal and grey matter damage(*37*, *38*).

**Figure 4.**
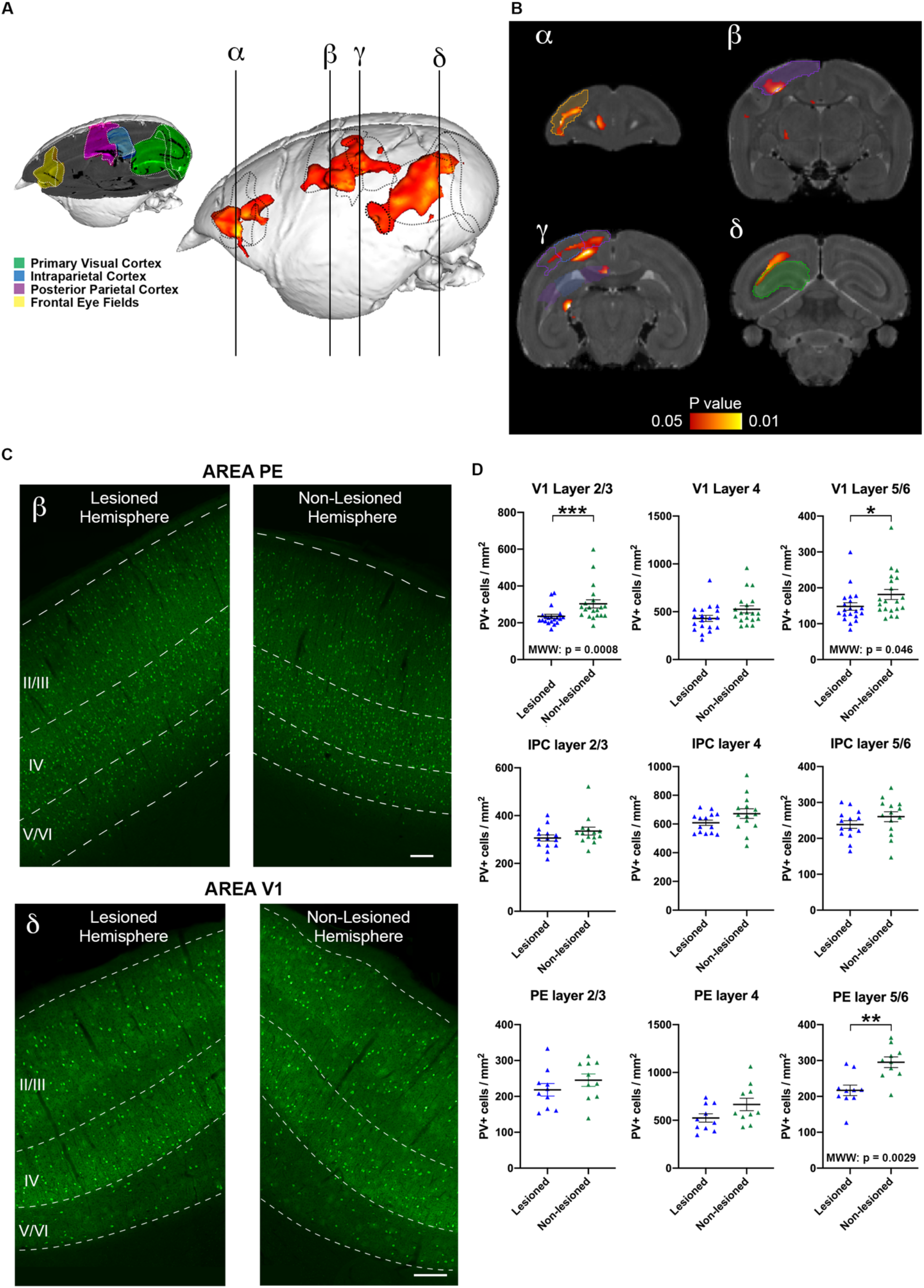
Microstructrual changes to the cortex following a unilateral MT lesion, as revealed by Diffusion Tractography Imaging (DTI) and parvalbumin (PV) fluorescent immunohistochemistry. **(A)** Voxel-based morphometry of fractional anisotropy (FA) maps. Yellow-red clusters represent significant hemispheric differences in FA, which was lower and observed in the ipsi-lesional primary visual cortex, intraparietal cortex, posterior parietal cortex and frontal eye fields (Left vs Right, n=5). Vertical lines (α, β, γ, δ) represent coronal sections in **(B)**. **(C, D)** PV+ interneurons were reduced in the lesioned hemisphere of areas PE and V1 (scale bar = 200μm). Scatter plots show significant reduction in PV neuronal density in infragranular layers of PE (p value = 0.0029) as well as supragranular layers (p value = 0.0008) and infragranular layers (p value = 0.046) of V1.

As we had previously demonstrated, there is a correlation between changes in FA and remodelling of the local circuitry(*39*), therefore we examined more closely the cellular neuroanatomy in the areas affected by the lesion of MT. Specifically, quantification of the calcium-binding protein, parvalbumin (PV) positive interneurons was undertaken as PV+ neurons have garnered significant interest as a component in the maturation of the neocortex and are functionally capable of amplifying circuit function(*40*–*42*). When compared to the non-lesioned hemisphere, PV+ cell density within the ipsilesional hemisphere was reduced in V1 (Mann-Whitney U test. Layers 2/3, p value = 0.0008; layers 5/6, p value = 0.046) and PE (Mann-Whitney U test. Layers 5/6, p value = 0.0029)**(Fig 4C, D)**. Although there was a reduction in of PV+ cells in the other layers of V1 and PE, as well as IPC in the lesion animals, these were not significant.

## Discussion

This study was conceived with the objective to determine how MT contributes to development of the dorsal stream. Further, an early-life unilateral lesion of MT would enable us to determine the lifelong behavioural deficits associated with loss of this central area of the dorsal stream and the impact this has on upstream and downstream cortices. Clinical studies which examined the consequence of losing V1 in early life vs adulthood demonstrated a marked difference in behavioural performance(*26*–*28*). Patient BI, whom suffered a bilateral V1 injury at nine days of age retains the ability to perceive colour and possesses significant conscious vision. When comparing the visual abilities of BI to an extensively studied blindsight patient, such as GY, who sustained an injury to V1 at the age of 8, BI possesses greater preservation of vision. As there is a strong correlation with the age of when an injury is acquired and the preservation of abilities that neuroplasticity affords, we wanted to explore the effects an MT lesion would have on visuomotor behaviour in a developing visual system. Additionally, we wanted to determine if there are behaviours and brain areas which critically depend on MT. From our results, it is clear that an early-life lesion of MT has a significant impact on visual system behaviour and architecture and provides evidence that area MT is crucial in the normal establishment of the dorsal stream network.

Seminal works studying the effect of lesions to MT on visual behaviour have unfortunately only looked at adult acquired loss. Newsome and colleagues provided awake behavioural evidence in the macaque that sensitivity to motion coherence is significantly disrupted following lesion to MT(*43*). Furthermore, MT is required to accurately track moving stimulus with smooth pursuit eye saccades. Visuomotor behaviour following bilateral MT lesion in adulthood has been examined in the macaque(*44*). The subjects were presented a battery of tests to probe reach and grasp behaviour in static and moving objects. Most notably, MT lesioned animals showed impairment in their ability to engage in a static reach and grasp task whereby the food reward is placed in a narrow slot so that the animal must only use the index and thumb for successful retrieval of the food reward. Closer examination of the retrieval mechanism employed by the adult MT-lesion animals showed they had difficulty in orientating their hand and typically had to grasp the food reward with their whole hand and were unable to exhibit fine grasps with their fingertips. The authors were unable to determine the impact the adult acquired bilateral MT loss would have on visuomotor behaviour in moving objects. The moving object retrieval task in their study consisted of a spinning bowl which would rotate at 240RPM. A banana pellet would be dropped in the spinning bowl and as such bounce around. All animals had difficulty with this task and no significant difference was observed between control and lesioned cohorts. In our study, our animals also showed perturbations in hand posture when engaging in the static task as seen with MT lesioned animals, exhibiting a larger MGA when reaching for food reward. This suggests that MT is required for prehension throughout life. However, it is possible that the deficits observed in this adult lesion model are due to the lesion extending beyond MT, as removal of underlying white matter was a component of the ablation and degeneration of the LGN observed, suggestive of damage to the optic radiation.

Clinically, there has only been one case of cerebral motion blindness described. Patient LM suffered a bilateral vascular insult to the temporal portions of her brain in adulthood, which included MT, as an adult with no injury evident in V1(*23*). Visual assessment of patient LM demonstrated that she had perturbations in reach and grasp behaviour, oculomotor scanning patterns and significant impairment in perceiving moving stimulus, although her ability to discriminate the colour and form of objects remained normal(*24*, *45*). While LM clearly had akinetopsia, she demonstrated an ability to reach and grasp static objects, and objects which moved up to speeds of 0.5m/sec (*46*). At speeds greater than 0.5m/sec, her performance greatly diminished and often executed her reach with greater velocity.

Our results share certain similarities with what has been observed in LM. Early-life MT lesioned marmosets also performed poorly when intercepting moving objects and displayed greater peak velocity and acceleration in their executed reach compared to controls during the static task. A notable difference between our study and LM’s capabilities is that LM performed remarkably well when reaching for objects that were static or slower than 0.5m/sec and her ability to intercept objects at low speeds was on par with that of controls. Clinically, there is evidence that even with static object retrieval, transcranial magnetic stimulation of MT interferes with fluid goal-directed reaching(*47*). Our early-life lesioned marmosets, as well as the macaques with adulthood lesions in MT in the Gattass and colleagues’ study, were unable to replicate the success LM possesses with static object retrievals. We believe LM’s ability to reach out to stationary and slow-moving stimuli could be a testament to her remarkable adaptive strategies in utilising the static visual cues available to her to estimate trajectories of slow-moving items and her hand relative to the stationary target object. As the marmosets in our study displayed impairment in retrieval of moving objects akin to patient LM, we postulate that the plasticity of the developing brain does not permit for recovery of visuomotor function in moving targets. As such MT serves a critical node for visuomotor behaviours which require information from moving stimulus throughout life.

Further dissection of the failed behavioural trials by each marmoset in the moving object task revealed that the lesioned cohort initiated the reach prematurely. This observed behaviour could be due to the animal’s inability to accurately detect the speed of the moving stimulus - akinetopsia, a deficit which was also observed in LM. It should be noted that motion is encoded in multiple cortical areas and not just MT(*48*, *49*). Therefore, the intrinsic ability of the brain to translate motion perception into a meaningful and precise goal-directed action relies on more than just sensitivity to motion coherence and further reinforces how integral MT is for object interaction, and the integral role it must place in the establishment of the associated networks during development.

Existing literature has highlighted the importance the medial intraparietal area (MIP) and parietal area (PE) serve in reaching actions and limb coordination(*50*, *51*). Considering we observed anatomical alterations specifically in these areas following an early-life lesion of MT, we suggest they are complicit in the perturbed reaching behaviour observed in both the static and moving object tasks. However, this does not explain why the lesioned marmosets demonstrate a similar kinematic profile as their control counterparts when intercepting moving targets. It is most likely, as marmosets do not have a refined hand use akin to macaques or chimpanzees(*52*, *53*), that the tendency to execute reach and grasp tasks on moving targets with maximum velocity is their natural teleological behaviour, especially for fast arboreal manoeuvres. Despite the comparable kinematics in the moving object task, ultimately the lesioned animals tended to prematurely reach and fail to collect moving object.

To the best of our knowledge, this is the first study examining the tuning properties of V1 neurons following an early-life injury to an extrastriate area. Previously, studies have examined the physiological properties of neurons in MT following a lesion of V1(*54*, *55*), revealing receptive field size and tuning properties were dramatically affected. However, MT responses remained robust and largely unaffected following V1 lesions sustained in early life(*56*). The only noticeable deficit was a reduced proportion of DS neurons. This supports the idea that the visual brain is largely able to execute contingency mechanisms during development to afford a higher level of functional recovery following the early-life loss of V1. This concept has also been observed clinically in subject BI, who following a bilateral early-life injury to V1 has significant preservation of vision, including sensitivity to moving stimuli(*27*). We wanted to observe the converse; if the tuning properties of V1 are vulnerable during development and extent of co-dependency between V1 and MT to develop DS or any other tuning property. In this study, we observed no dramatic perturbations in the tuning properties of V1 neurons. We did observe a lower proportion of DS neurons, which parallels previous findings for MT following an early-life lesion of V1(*56*). Modifications of V1 tuning properties, such as DS or even orientation selectivity has been reported before(*57*–*59*) but none specifically in relation to V1-MT circuitry. This warrants further investigation to determine if the MT-V1 projection serves a role in early life to tune DS within V1 neurons.

As PV interneurons form the majority of GABAergic interneurons in V1(*60*), we sought to determine if there were disruptions to the local circuits by quantifying PV+ interneurons. Further, it is well documented that the local interneuron circuitry is more susceptible during development than in adulthood(*61*). MT has considerable reciprocal connectivity with layers 3C and 6 of V1(*5*),and we revealed a reduced PV+ cell density within both supra and infragranular layers. PV+ neurons in V1 have a diversity of feature-specific visual responses that include sharp orientation and DS, small receptive fields, and bandpass spatial frequency tuning(*62*). These results suggest that subsets of parvalbumin interneurons are components of specific cortical networks, and that perisomatic inhibition contributes to the generation of precise response properties. Ablation of PV in V1 interneurons has previously been observed to decrease DS properties within V1(*63*). Therefore, it is conceivable that the reduced PV population in V1 following the early-life lesion of MT is the underlying basis to our observation of a lower proportion of DS neurons in V1.

The implications of this current study extend our understanding of the central role area MT occupies in the early establishment of the dorsal stream and associated behaviours, and its role as an anchor in the developing visual cortex. Further, our observation of a lower proportion of DS neurons within V1 could suggest a level of dependency on MT in early life in the tuning and appropriate maturation of V1 neurons. While marmosets lack fine motor skills, they have demonstrated their ability to use tools(*64*) and have the circuitry for accurate reach and grasp behaviours. Teleologically, the early establishment of the dorsal stream network provides primates the capacity to process vision-for-action, including accurate reaching and grasping behaviour, which is integral to their survival.

## Materials and Methods

### Subjects

Five New World marmoset monkeys (*Callithrix jacchus*) received a mechanical ablation of the middle temporal (MT) area at postnatal day (PD) 14 **(Table 1)**. Following surgical ablation, the animals were allowed 12 months of recovery before undergoing training for visual behaviour experimentation. Three aged-matched animals were used as controls for behavioural experiments (Table 1). All experiments were conducted in accordance with the Australian Code of Practice for the Care and Use of Animals for Scientific Purposes. All procedures were approved by the Monash University Animal Ethics Committee, which also monitored the health and wellbeing of the animals throughout the experiments.

**Table 1.** Marmosets employed in current study. Prefixes in animal ID: M=male; F=female.

| *Animal ID* | Neonatal unilateral MT Ablation | MRI/DTI | Reach & Grasp Behaviour | V1 Electrophysiology | Immuno-histochemistry |
| --- | --- | --- | --- | --- | --- |
| *M1695* | X | X |  | X |  |
| *F1696* | X | X |  | X |  |
| *F1992* | X | X |  |  | X |
| *F2019* | X | X | X |  | X |
| *F2020* | X | X | X |  | X |
| *F1565* |  |  | X |  |  |
| *F1575* |  |  | X |  |  |
| *M2086* |  |  | X |  |  |

#### Mechanical ablation of area MT

Neonatal marmosets (PD 14; n=5) were anaesthetised with isoflurane (1-2% in medical oxygen at 0.5L.min^−1^) and placed into a custom built non-ferromagnetic stereotaxic frame to facilitate T2-weighted MRI to allow for demarcation and targeted ablation of area MT **(Fig. 1B)**. Full details of preparation of the animal and how to localise areas of interest with MRI guidance have been described previously(*65*). Scans were exported as DICOM files and the brain images were visualised on the Horos v3.3.5 software (https://horosproject.org). Once MT was visualised, ablations were performed using a biopsy punch (diameter = 3.0 mm). Following ablation of cerebral tissue, the cavity was filled with Gelfoam, the cranium was reconstructed and the scalp was sutured. Animals would receive post-surgical antibiotic and analgesic medication and then allowed a minimum of 12 months recovery. to allow for plastic reorganisation of the brain.

#### Longitudinal MR imaging

During the animal’s recovery, structural T2 images were acquired on a 9.4T small-bore animal scanner (Agilent) 6 weeks and 36 weeks post lesion to identify the extent of the lesion **(Fig. 1B)**. The parameters for the T2 scan are: MRI structural images comprised T2 weighted images (9.4 Tesla 18 cm bore MRI scanner (Bruker); 2D-RARE sequence, TR=12000 ms, TE=48 ms, Matrix size=200×180, FOV= 40×36mm2, number of slices = 80, slice thickness=0.4mm, RARE factor = 8, axial plane, and 4 averages; scan time 30 minutes). Additionally, at the same time points, diffusion tensor imaging (DTI) was acquired for analysis of changes in microstructure in visual cortical tissue following injury.DTI was acquired in the axial orientation using a six-shot spinecho echo planar imaging (EPI)-based DTI sequence. Scan parameters included: TR/TE of 3,000/31 ms, FOV of 38.4 × 38.4 mm^2^, data matrix of 96 × 96, and 50 slices with thickness of 0.4 mm. Diffusion encoding gradients were applied in 30 directions, with a b value of 800, gradient duration of 5 ms, and separation of 18 ms. A reference scan with inverted phase and readout gradients was acquired for phase correction to reduce EPI and DTI-specific artefacts. The average of the reference and the original scan also increased the signal-to-noise ratio (SNR). The DTI sequence was repeated four times and saved separately with the aim of further increasing SNR and generating backup data in case any unexpected motion artifacts occurred during the relatively long scan. The whole DTI scan took ∼80 min.

Diffusion weighted data were pre-processed and analysed using tools from different software packages that work well with the small sized marmoset brain. MriCron, dcm2nii script (http://people.cas.sc.edu/rorden/mricron/dcm2nii.html) was used for conversion of DiCOM to nifti files; averaging, orientation of different DWI repetitions and corrections for movement distortions was scripted using FSL 5.0.9 software library of the Centre for Functional Magnetic Resonance Imaging of the Brain (FMRIB, University of Oxford, UK, www.fmrib.ox.ac.uk/fsl). Binary brain masks were generated using Brainsuite 15c (www.brainsuite.org). Diffusion tensors and derived fractional anisotropy maps (FA) were calculated using MRtrix3 (www.mrtrix.org). (*66*)

#### Visual Behaviour Training

Visual behaviour experiments commenced following a minimum of 12 months recovery from the MT ablative surgery. Detailed stepwise training and habituation protocols to allow for implementing reach and grasp behavioural experiments in the marmoset have been previously described(*32*).

Animals were first trained to enter a custom fabricated behaviour training box (Monash Instrumentation Development Facility). Food rewards are given to positively encourage participation in desired behaviours. Following habituation with the behaviour box, the animal is transferred to an adjacent room where the behavioural experimentation is conducted. Food rewards are once again given to habituate the animal with the room.

Following complete habituation, the animal’s ability in goal orientated reach and grasp of static objects was examined whereby the animal is presented with a target object in the form of a food reward (piece of fruit). The food reward was placed in one of two positions on a pedestal, both of which are equidistant to an aperture in the behaviour box **(Fig. 2A-D)**. These two positions could favour either the left or right hand. To determine the animals’ hand preshaping behaviour and if the size of the target object would have a significant effect on performance, the food reward was cut into cuboids where the largest face would be ~5×5mm or ~10×10mm. The precise size of the food reward was qualified during the video analysis of each trial with in- built tools in Tracker software. Before commencement of a static task trial, a blind is placed in front of the transport box and its apertures so the animal could not prime its hand in anticipation of the position of the reward. Each static object retrieval session consisted of 15 - 20 trials. The proportion of trials of small vs large object presentation was equal.

To assess the animal’s ability in reaching and grasping a moving object, a single sized food reward (cuboid, ~10×10mm face) was placed on the edge of a custom-built rotating turntable (ø=14cm, Monash University Instrumentation Development Facility)**(Fig. 2b, D)**. The centre of the turntable was aligned with the centre of the transport which was equidistant to both right and left apertures. Objects were always placed in the same spot of the turntable across all trials. Shutters were used to occlude the reaching apertures to control the use of hand (i.e, the animal would use its right hand when the right aperture was unblocked and vice versa). The shutter was removed, and the trial commenced once the food reward had undergone one revolution to ensure that the turntable can accelerate to the desired speed for the task. Animals were given a 10-second response time for each trial. Animals were allowed a maximum of three attempts within each trial for it to be considered successful. Background white noise was played throughout the sessions to minimise any impact the noise produced by the turntable would have in the animal’s performance.

Three independent variables were incorporated to the turntable task; 1) Six different revolution speeds, which ranges from 10 to 60 revolutions per minute (RPM), 2) Direction of the revolution (clockwise/anti-clockwise), and 3) Hand used to retrieve the object **(Table 2)**. This resulted in 24 different conditions that would be presented during a training session. The order of the trial conditions randomised across sessions and a single condition (eg. object moving at a speed of 30RPM in a clockwise direction with the aperture opened for the left hand) did not repeat in the same session.

**Table 2.** Independent variables employed in moving reach and grasp task

| Independent Variable | Description |
| --- | --- |
| Speed | 10, 20, 30, 40, 50, 60 RPM |
| Direction | clockwise or anti-clockwise |
| Hand Use | Aperture blocked to allow for only the left or right hand for object retrieval |

#### Analysis of visual behaviour performance

Trials for the static object task were binned into three main categories; a successful trial was defined as a single coordinated motor action to retrieve the object. Corrected trials occurred when animals required fewer than three corrections in either their reach or grasp to retrieve the object successfully. Failure to meet the conditions of the two previous bins, or not completing the action within the given time frame, was assigned to the failed trial category. Trials for the moving object task were binned into two main categories; a successful trial when the animal retrieved the rotating object within three attempts, a failed trial when more than three attempts were required for retrieval or the object was dislocated from the initial position. Trials with no response were considered invalid and discarded.

All trials were recorded using “GoPro HERO4 Black” camera (1280×790-pixel resolution, 120 frames/sec, narrow view) from the transverse plane perpendicular to the plane of motion (i.e. top-down view) and videos captured were analysed offline. The same investigator (C-K.C) performed and analyzed all videos offline manually in a blinded manner to determine the performance and kinematics of each trial. The performance as expressed by error rate was determined by dividing the number of failed trials over the total number of trials within a given session. The overall performance of a given task in each animal was determined with permutation testing of the performance of all sessions.

Video files were transferred to an open source software Tracker v4.93 (physlets.org/tracker/) designed to perform a 2D kinematic analysis. A reference scale of known size remained in the field of view and was positioned at the height of the reaching movements to eliminate any perspective errors. The edges of the reaching platform served as reference points for X and Y axes to track movement trajectories across each trial. Three metrics were measured in the video analysis. The maximum grip aperture (MGA) between the animal’s thumb and index finger when the animal pre-shapes its hand in acquiring an object, the velocity at which the animal would move to grasp an object as well as the acceleration. Additionally, the junctions between the nail and digit of the index finger and thumb were used as reference points to measure the Euclidean distance over time. Movement duration was defined as the time elapsed from the moment the thumb reference point was visible outside of the reaching aperture in the box to the moment when either the index finger or thumb made contact with piece of fruit (identified by it moving for at least three consecutive frames). The length of the food reward in each trial was measured against the reference scale and plotted against the MGA of the trial using Prism version 8.4.2 to determine if the animal exhibited any hand preshaping behaviour. All failed trials were further sorted to determine whether the animals exhibited a premature or delayed reach while attempting to retrieve the food reward. The angle between the final hand position over the turntable and the food reward was then measured on Tracker v4.93 to determine the accuracy of targeting the object **(Fig. 2I)**. Mean angulation was calculated in each session for each animal to reveal overall angulation of failed trials across all sessions. *P*≤ 0.05 were considered to be statistically significant. All data are presented as mean ± standard error of the mean SEM).

#### Electrophysiological recordings

Animals M1695 and F1696 underwent a single recording session under deep anaesthesia. The detailed methodology of how the animals were prepared for electrophysiology is described in Bourne & Rosa(*67*). In brief, animals underwent a tracheotomy and two craniotomy procedures with anaesthesia being maintained with a continuous intravenous infusion of a mixture of pancuronium bromide (0.1 mg.kg-1.h^−1^), sufentanil (6 μg.kg-1.h^−1^), and dexamethasone (0.4 mg.kg-1.h^−1^), in a saline–glucose solution. During the recordings, animals were also ventilated with nitrous oxide and oxygen (7:3).

The electrophysiological experiments consisted of two phases. Firstly, the scotoma of each animal was mapped by determination of MT complex receptive fields along the perimeter of the lesion core. Following identification of the scotoma, recordings were conducted in V1 within and outside the lesion projection zone. Recordings were performed with tungsten microelectrodes (~ 1MΩ). Insertion of the electrodes to our eccentricity of interest was guided by stereotaxic and topographical data described in previous studies(*5*, *68*). Amplification and filtering were achieved via an AM Systems model 1800 microelectrode alternating current amplifier. Stimuli presentation and methods for quantification of tuning for orientation and direction as well as spatial and temporal frequency within V1 was performed as per previous studies(*35*, *56*, *69*).

#### Histology and immunohistochemical tissue processing

Animals were euthanised with an overdose of sodium pentobarbitone (>100mg/kg) then transcardially perfused with 10mM PBS that has been supplemented with heparin (50 IU/ml of PBS), followed by 4% paraformaldehyde in 10mM PBS. Brains were post-fixed in 4% paraformaldehyde then dehydrated in serial solutions of PBS-sucrose before being snap frozen in 2-methylbutane that has been chilled to −50°C and stored in a −80°C freezer until cryosectioning.

Brain samples were cryosectioned in the coronal plane at 50um, divided into five series and stored free-floating in a cryoprotective solution (as outlined in previous work(*3*)).

Half a series was stained for myelin via silver impregnation(*70*) to demarcate area MT and to qualify the size of the lesion and spatially, the extent of MT which was ablated **(Fig. 1C)**. Free-floating sections were labelled with GABAergic interneuronal marker parvalbumin (PV; Swant, PV27 1:3000).

#### Microscopy and digital image processing

Brain sections were imaged with an Axio Imager Z1 microscope (Zeiss). Images were obtained with a Zeiss Axiocam HRm digital camera using Axiovision software (v. 4.8.1.0) at a resolution of 1024 by 1024 or 2048 by 2048 pixels and saved in ZVI and exported to TIFF format. The objectives used were Zeiss EC-Plan Neofluar 5×0.16, #420330-9901, EC-Plan Neofluar 10×0.3, #420340-9901. Filter sets used for visualizing immunolabelled cells were Zeiss 38 HE eGFP # 489038-9901-000.

Images used for cell density quantification were taken with the 10x objective. Stitching of images and adjustments to contrast and brightness was performed using Adobe Photoshop 2020 v21.0.1 or Zeiss MosaiX software. The contours, boundaries, and line art of all figures were drawn using Adobe Illustrator 2020 v24.0.1.

Quantification of interneuronal density was conducted as per previous work(*30*). Each area or interest had analysis of PV cell density within the supragranular layers 2 and 3, granular layer 4 or infragranular layers 5 and 6. For each animal, 6 sections randomly selected across the anterior-posterior axis across our areas of interest. The regions that were sampled are V1, the areas around the intraparietal area which consists of areas AIP, LIP & MIP and area PE which is part of the posterior parietal cortex. Analysis of photomontages of CB and PV IR cells was conducted using Fiji image software(*71*). The same investigator (W.C.K) performed all quantifications in a blinded manner.

#### Statistical analysis

Assessment of the behavioural performance of an animal was determined through their error rate in each session. The error rate, as well as the frequency of successful retrieval on the first attempt was examined using a non-parametric permutation test on Microsoft Excel version 16.62. The kinematic analysis of reach and grasp behaviours which include MGA, velocity, acceleration and angulation was also examined using a non-parametric permutation test on Microsoft Excel version 16.62. Within the permutation test, data were repeatedly resampled (5000 times) to determine the difference between cohorts. *P*≤ 0.05 were considered to be statistically significant.

To determine if sizes of the food reward have any influence on the performance, an ANCOVA analysis was performed on Prism version 8.4.2 to compare the error rate of large and small sizes across sessions. A linear regression was also performed on Prism version 8.4.2 to determine the hand preshaping behaviour against food lengths. *P*≤ 0.05 were considered to be statistically significant.

Statistical analysis for PV+ cell density as presented as mean ± standard error of the mean was carried out using Prism version 8.4.2 software. Comparison between the ipsi-lesional and contra-lesional V1 were examined by a non-parametric Mann-Whitney U test. *P*≤ 0.05 were considered to be statistically significant.

For voxel-based morphometry statistical analysis of FA maps a general linear model following a two-sample t-test was used comparing left (lesioned) and right (control) hemispheres (SPM12, The Wellcome Trust Centre for Neuroimaging, UCL, UK) as follows. FA maps from all MT-lesion animals were registered to a marmoset brain template (http://brainatlas.brain.riken.jp/marmoset/. BSI-Neuroinformatics, Riken Institute, Japan) using the normalized mutual information function (7-degree B-spline). Resulting images were smoothed with 1 mm isotropic Gaussian kernel. Subsequently, images were flipped and registered again to original unflipped images. Significance level was P<0.05 corrected for the familywise error rate (FWE) with and extended threshold of 1 voxel. Z-scores converted to P values were displayed on the Brain/MINDS 3D digital marmoset atlas(*66*) using Mango 4.1 software from the Research Imaging Institute, University of Texas Health Centre at San Antonio (ric.uthscsa.edu/mango).

## Acknowledgments

The authors would like to acknowledge and thank D.A. Leopold who read through earlier versions of the manuscript and M.J. de Souza for technical assistance and Qi Zhu Wu for MRI acquisition and DTI preprocessing.

## Author Contributions

W.C.K. and J.A.B. conceived and designed the research. W.C.K., C-K.C., H-H.Y., I-C.M. and J.A.B. designed the experiments. W.C.K., I-C.M., D.M.F, J.A.B. performed the MT ablation surgeries. C-K.C., W.C.K., and D.M.F performed the behavioural experiments. C-K.C and D.M.F analyzed the behavioural data. W.C.K., I-C.M., H-H.Y., J-H.L., J.A.B performed the electrophysiology experiments. I-C.M. analyzed the MRI data. W.C.K performed all tissue processing, immunolabelling and cell counts. W.C.K., C-K.C., H-H.Y. and J.A.B. wrote the paper.

## Data and materials availability

All data that supports the findings of this study are available from the authors on reasonable request. See author contributions for specific datasets.

