## Supplementary figures and images for "Visual Cortical Area MT is Required for Development of the Dorsal Stream and Associated Visuomotor Behaviours"

### Supplementary Figure 1

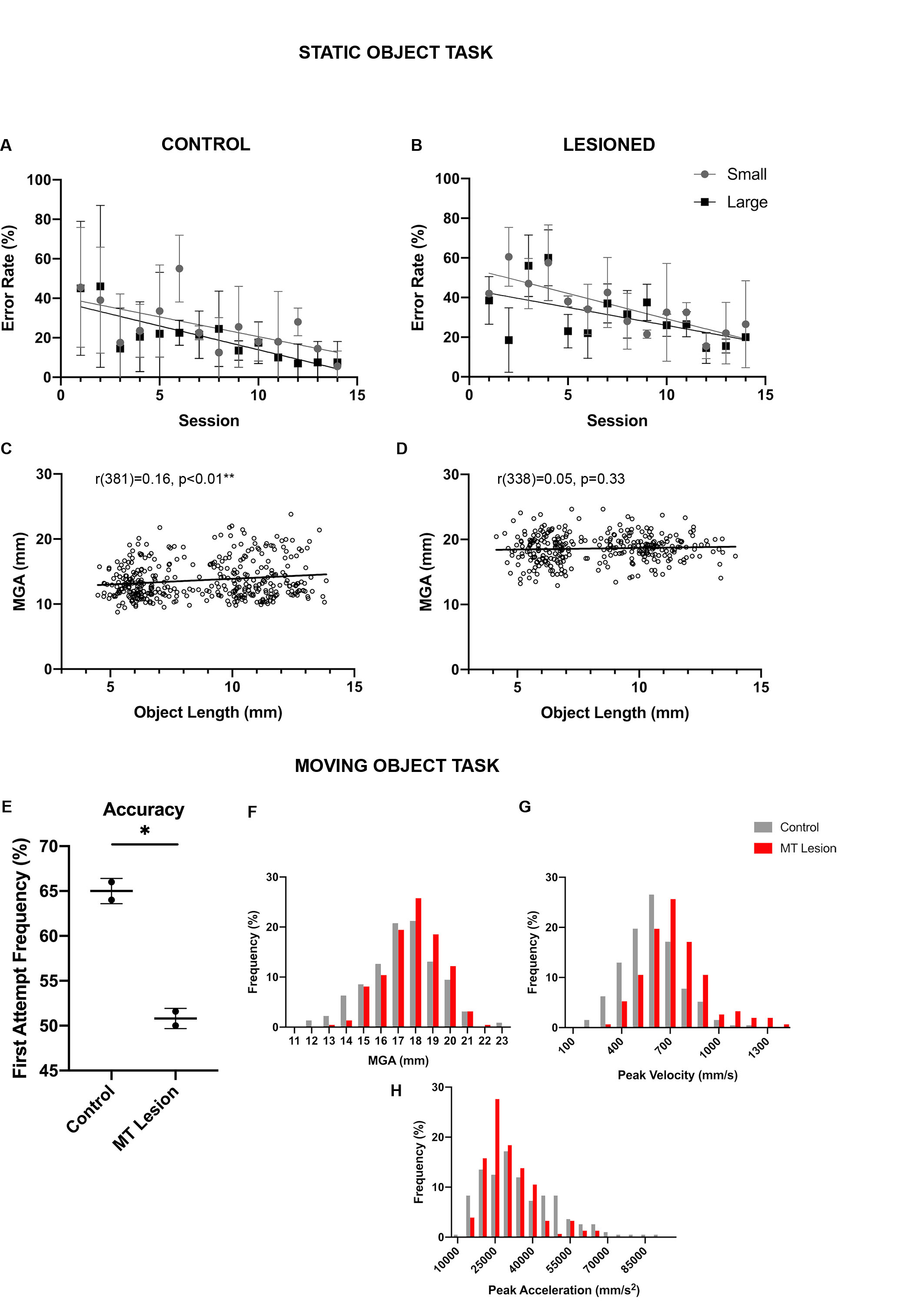
